# Scaling up by looking closer: extending the range of moss-Nostoc symbioses deeper into central Europe and to new moss hosts in the forest canopy

**DOI:** 10.64898/2025.12.22.695964

**Authors:** Feulner Martin, Ulla Spieleder, Frederic J. Pascual Dose, Jona Motzkus, Daniel C. Thomas

**Affiliations:** University of Bayreuth, Bayreuth Center for Ecology and Environmental Research (BayCEER); University of Bayreuth, Bayreuth Center of Ecology and Environmental Research (BayCEER); University of Goettingen, Program for Biodiversity: Ecology, Evolution, and Conservation (BEEC)

**Keywords:** *Nostoc*, symbiosis, central Europe, forest, moss

## Abstract

Since their re-discovery, Moss-cyanobacterial symbioses are of increasing interest, especially *Nostoc-*feather-moss symbioses. Efforts to document *Nostoc*-feather-moss symbioses have historically been centered in Scandinavia and the North American Boreal forest zone. Herein we report the results of several opportunistic surveys investigating the occurrence of *Nostoc-*moss symbioses in the region of Franconia (Germany). Using culture- and microscopy-based methods, we confirm the occurrence of such symbioses in our region. Two new moss host species of the epiphytic family Orthotrichaceae, *Orthotrichum* and *Zygodon*, are also documented, further extending the range of known moss hosts for *Nostoc-*like organisms outside the traditional feather moss (Hypnales) and *Sphagnum* clades. Full length 16S sequence alignments placed the moss-associated *Nostoc-*like organisms throughout our Nostocaceae phylogeny, with no obvious single symbiotic clade. We note the presence of multiple alternate nifH genes within the genomes of single *Nostoc-*like organisms, possibly enabling diazotrophy under variable ecological conditions. We also present evidence for possible endophytic colonization by cyanobacteria of stems of mosses.

## Introduction

Close associations between *Nostoc* cyanobacteria and mosses have been noted by naturalists for over a century. Among the first described species of *Nostoc* (sensu latu, Nostoc Vaucher ex Bornet and Flahault 1886) were specimens collected from mosses (*Nostoc muscorum* Agardh ex Bornet et Flahault 1888, syn. *Desmonostoc muscorum* (*Hrouzek et al., 2013*). Researchers throughout the twentieth century continued to note frequent associations of *Nostoc*-like organisms and mosses (division Bryophyta) (Meeks, 1990). The potential importance of these associations at larger scale in temperate and arctic terrestrial ecosystems was predicted and quantified in multiple early studies (Dodds et al., 1995). While all mosses are known to host cyanobacterial communities (Groß et al., 2024; Rousk, 2022), especially *Sphagnum* mosses (Rousk et al., 2015), interest in bryophyte-*Nostoc* associations grew rapidly following the documentation of a widespread symbiosis between a *Nostoc* sp. and a feather moss *Pleurozium schreberi* in boreal forests (DeLuca et al., 2002). The super-abundance of *Pleurozium* and other feather mosses (order Hypnales) in the large northern forest zones implied immense importance for global-scale N and C budgets. Since these findings, research on feather-moss/*Nostoc* symbioses has accelerated and studies have employed increasingly sophisticated means of detection and quantification of nitrogen-fixation and nutrient transfer (Alvarenga et al., 2024; Arróniz-Crespo et al., 2022; Bay et al., 2013; Stuart et al., 2020).

Moss-cyanobacterial mutualisms appear to strongly contribute to the primary production of boreal forests and therefore contributes to ecosystem services (timber production) and sustainability (reduced fertilizer inputs). Through associations with nitrogen-fixing microorganisms, mosses can provide large inputs of nitrogen for ecosystems. In boreal forests, up to 50% of the nitrogen input may come from cyanobacteria associated with mosses (Rousk & Michelsen, 2017). In peatland cyanobacteria can contribute more than 35% of the nitrogen required by mosses like Sphagnum, helping maintain the productivity and stability of these carbon-rich environments (Berg et al., 2013). In harsher climates, such as tundra regions (reviewed by Lindow et al. (2013)) or tree-canopy microclimates (Lindo & Whiteley, 2011), cyanobacteria associated with mosses can account for the vast majority of the nitrogen-fixation necessary for observed rates of net primary production. Variation in the community of nitrogen-fixing bacteria -- and especially cyanobacteria – appears to be explained by both geography (Groß et al., 2024) and the host specificity (Ininbergs et al., 2011).

Following classical, broad definitions of symbiosis as any long-term, physically-proximal biotic interaction (Tipton et al. 2019), symbioses between feather mosses and Nostoc-like organisms are generally considered to be “loose”, mutually-facultative or commensal exosymbioses. *Nostoc*-like organisms appear to prefer to epiphytically inhabit the spaces in protected leaflet-axils along moss caulids (“stems”), or in incurved regions of leaflet-tissue and other protected areas (DeLuca et al., 2002; Solheim et al., 2004), suggesting at least a structural benefit for *Nostoc*-like organisms when associating with moss hosts. Indeed, the main advantage for the cyanobacterium may be a buffered environment on the moss, as the hygroscopic nature of mosses may create sheltered microenvironments even during times of high radiative exposure (Maeng et al., 2025) or drought (Li et al., 2024) for their resident microbiome.

Benefits from the moss-*Nostoc* symbioses for the moss host are presumed, as chemo-attraction and induction of mobile hormogonia phase by feather-mosses in Nostoc symbionts have been clearly documented (Bay et al., 2013), at an energetic cost to the host moss (Sprent & Meeks, 2013). Nutrient exchanges are hypothesized, involving a transfer of fixed organic nitrogen in the form of fixed, organic nitrogen from heterocystous Nostoc colonies to the host plant (Arróniz-Crespo et al., 2022; Bay et al., 2013), perhaps in exchange for organic carbon- and sulfur-containing materials from the moss host, though the latter has only recently been directly reported (Stuart et al., 2020). Symbioses vary seasonally and daily (Warshan et al., 2016) in their levels of nitrogen-fixing activity and microbial abundance, due to climatic constraints on microbial metabolism (Gentili et al., 2005) and N-availability at both host (Renaudin et al., 2022) and ecosystem scales (Cusack et al., 2009; Zheng et al., 2019).

Traditionally, *Nostoc*-moss symbioses have been studied most intensively in Scandinavian and North American regions (Alvarenga & Rousk, 2022), but they have been observed in many extraordinary sites throughout the world, including retreating glacial fields of Tierra del Fuego (Chile) (Arróniz-Crespo et al., 2014), Andean tropical montane cloud forest (Peru) (Permin et al., 2022) and Mount Fuji (Japan) (Kubota et al., 2023). Arboreal mosses are increasingly of interest, and have been observed to be hotspots for moss-cyanobacterial symbioses (Lindo & Whiteley, 2011). Despite its proximity to Scandinavia and despite many shared moss species and sites of similar environmental conditions, direct observations of the symbiosis of feather mosses and *Nostoc* (or *Nostoc-*like organisms) are rare from the Central European region. To our knowledge the *Nostoc* symbiosis has been observed in German mosses only once, as one member out of many nitrogen-fixing bacteria associated with log-dwelling moss species, using nifH metabarcoding (Groß et al., 2024). Traditional “model” species of feather-moss involved in *Nostoc* symbioses like *Hylocomium* (Fig. 1) or *Pleurozium* have to date received very little attention in central Europe, despite the prevalence of these moss species in montane forests of central Europe. Particularly likely to host moss-*Nostoc* symbioses are spruce- or pine-dominated forests, as *Pleurozium schreberi* or *Hylocomium splendens* are among the dominant floor mosses in these forests (Ellenberg, 1988).

**Figure 1:**
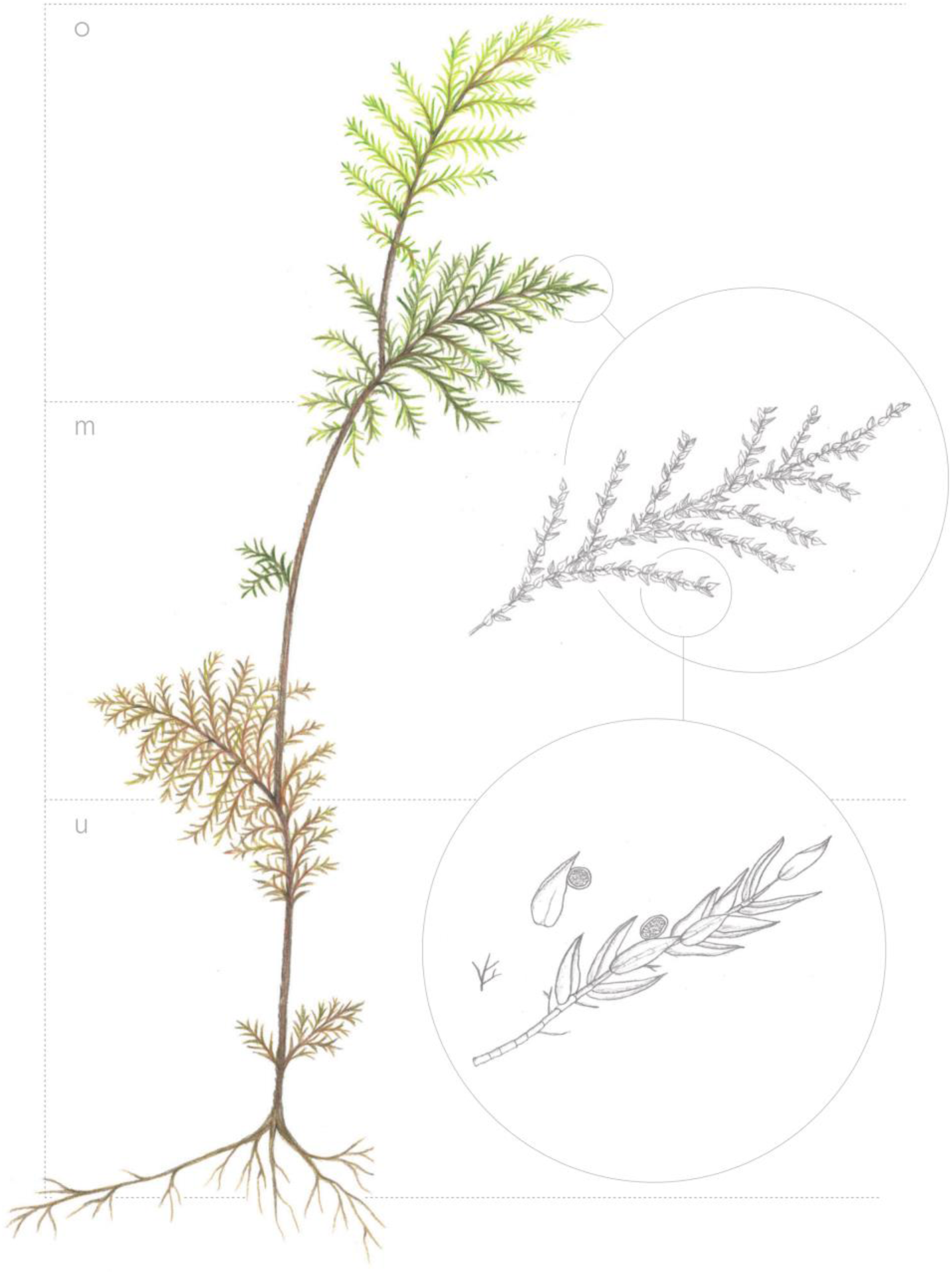
Hylocomium splendens with colonies of Nostoc ((c) J. Motzkus 2024)

Here we sought to confirm the presence of the moss-*Nostoc* symbioses in the region of Franconia, Germany, and to further explore the known host range for the symbiosis. We examined mosses from the Fichtel mountains, the Frankenjura, and adjacent areas, concurrent with on-going moss diversity inventories and monitoring efforts the region. Special attention was given to arboreal epiphytic mosses, especially in the family Orthotrichaceae. despite never having been documented as hosts for the symbiosis. Members of Orthotrichaecae are best known as epiphytic mosses from tree bark (Fialová et al., 2023), in our region especially on broad-leaved trees with many species. We hypothesized them as likely candidates to host the symbiosis, given the presumed low nitrogen availability and dynamic microclimate of the canopy setting. We utilized a microscopy- and culture-based approach for detection and description of *Nostoc*-like organisms, which we hoped would enable precise imaging and taxonomic placement of symbionts, using full-length 16S and partial nifH PCR amplicon sequence data.

## Methods

### Field

Mosses from multiple sites through the region of upper and middle Franconia (Germany) were opportunistically sampled for *Nostoc*-like organisms, as part of ongoing moss diversity inventories in the region. Moss specimens were collected when visual inspection with a 10x lupe indicated possible cyanobacterial colonies, in the form of blue discoloration. Moss samples were collected from multiple different locations in the Fichtel Mountains, the Frankenjura, and other sites at elevations between 350 and 800 m above sea level. Sample collections were carried out as part of other ecological surveys which took place during the years 2023 - 2025, see Table 1 for details. Each moss sample consisted of at least one healthy gametophyte individual (all continuous tissue from the rhizoids to the shoot). Samples were stored in plastic bags and kept in the dark at 4 °C before further use.

**Table 1.**
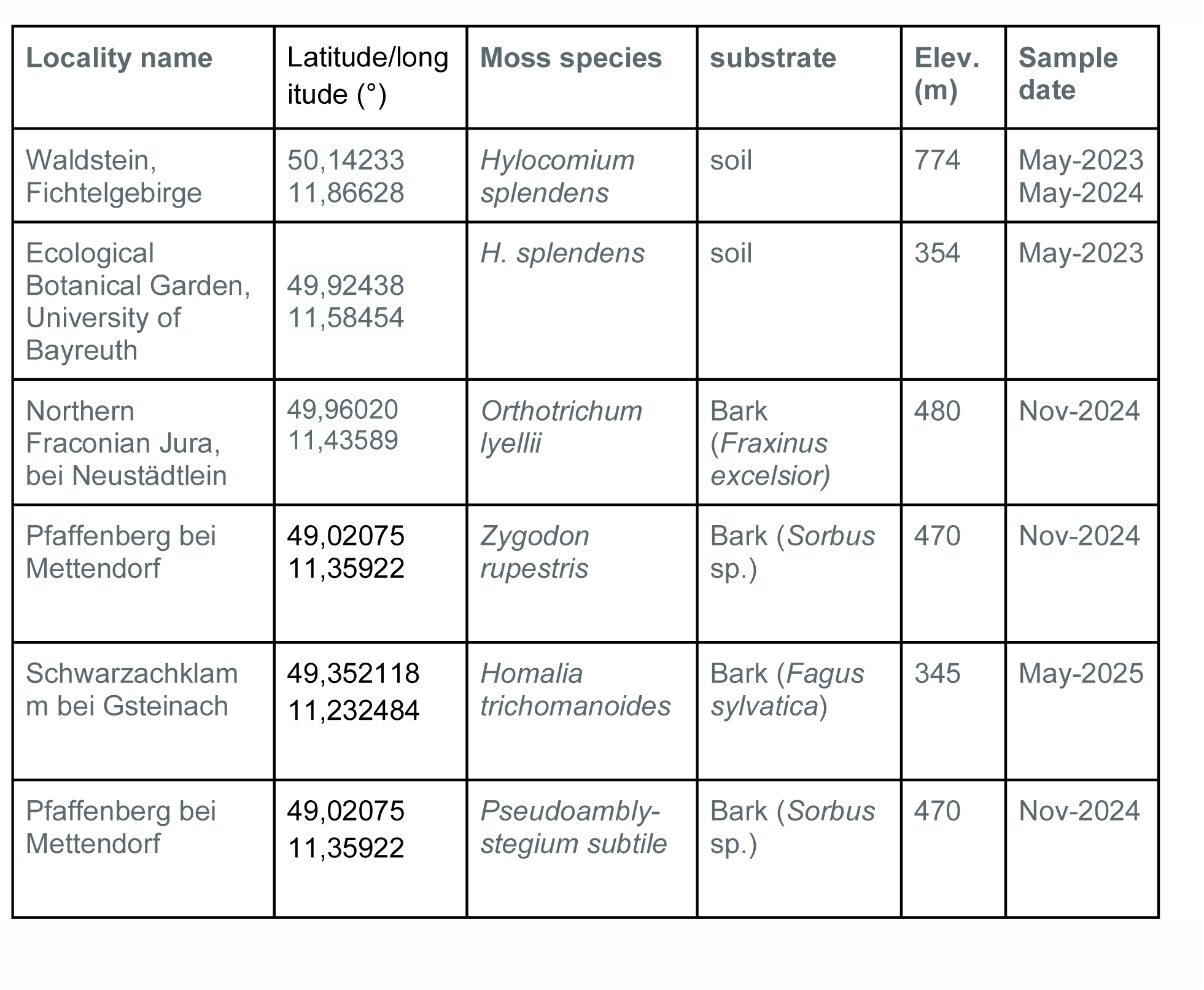
Sampled locations.

| Locality name | Latitude/longitude (°) | Moss species | substrate | Elev. (m) | Sample date |
| --- | --- | --- | --- | --- | --- |
| Waldstein, Fichtelgebirge | 50,14233<br>11,86628 | *Hylocomium splendens* | soil | 774 | May-2023<br>May-2024 |
| Ecological Botanical Garden, University of Bayreuth | 49,92438<br>11,58454 | *H. splendens* | soil | 354 | May-2023 |
| Northern Franconian Jura, bei Neustädtlein | 49,96020<br>11,43589 | *Orthotrichum lyellii* | Bark (*Fraxinus excelsior*) | 480 | Nov-2024 |
| Pfaffenberg bei Mettendorf | 49,02075<br>11,35922 | *Zygodon rupestris* | Bark (*Sorbus* sp.) | 470 | Nov-2024 |
| Schwarzachklamm bei Gsteinach | 49,352118<br>11,232484 | *Homalia trichomanoides* | Bark (*Fagus sylvatica*) | 345 | May-2025 |
| Pfaffenberg bei Mettendorf | 49,02075<br>11,35922 | *Pseudoamblystegium subtile* | Bark (*Sorbus* sp.) | 470 | Nov-2024 |

### Microbiology

#### Overview

All moss samples were screened microscopically for colonization by *Nostoc*-like species, first with classical light microscopy, followed by additional wide-field epifluorescence microscopy in specimens where colonies were observed. Following microscopy, culture of all observed spherical *Nostoc-*like colonies was attempted, and amplification of NifH and 16S rRNA marker genes, using colony PCRs. Whole community DNA (moss and microbiome) was extracted for some samples for PCR-based detection of nifH in moss tissues. In some cases, surface-sterilization was performed to detect the possibility of endophytic colonization by cyanobacteria of moss tissues.

#### Microscopy

Preliminary surveys for the presence of *Nostoc* were conducted with standard light microscopy, first using binocular stereoscopes (Wild HeerBrug M-series) and later using Zeiss compound microscopes. For comparison, some specimens which were surveyed for *Nostoc* colonies using light microscope and that were found not to host visible colonies were then subjected to DNA purification and to whole-community PCR amplification of nifH genes in PCR tests. Sectioning was done using sections of one centimeter from the top (o), middle (m) and bottom (u) of caulids (Fig. 1), for comparison of relative abundances of nifH abundances among different positions. Samples with possible *Nostoc* colonies were then subjected to further investigation with epifluorescence microscopy.

Moss tissues and bacterial cultures were also observed under an epifluorescence microscope (Revolve 2, ECHO, San Diego, USA), using Revolve Software 4.1.0, app version v7.1.0. With epifluorescence microscopy, cyanobacteria can be easily more discerned from eukaryotic microalgae and plant cells, as cyanobacteria present a distinct autofluorescent “fingerprint” characterized by phycobilisome-mediated fluorescence (Grigoryeva, 2020; Wojtasiewicz & Stoń-Egiert, 2016). As such, cyanobacterial cells can be identified using the standard “Cy5” filter setup (CHROMA 49009, ex: 610-650 nm, em: 660-735 nm), which detects C-phycocyanins (phycocyanins found only in cyanobacteria) with an excitation peak at around 625 nm (em: ≥650 nm) (Grigoryeva, 2020; Poniedziałek et al., 2017). In contrast, chlorophyll a fluorescence shows an excitation peak at 458 nm and ∼680 nm (em: ∼680 nm). As such, algal and plant cells — which contain significantly higher concentrations of chl a and no C-phycocyanin; (Wojtasiewicz & Stoń-Egiert, 2016) — can be detected with the “FITC/Cy2” filter set (CHROMA 49002, ex: 450-490 nm, em: 500-550 nm), which permits some detection of emissions from Chl a and select other embryophyta plant cell components, especially in cell walls (Donaldson, 2020; Tobimatsu et al., 2013). Phycoerythrin, which is present in varying concentrations in different cyanobacterial species, can also be detected using the “TXRED” filter (CHROMA 49008, ex: 540-580, em: 590-665 nm).

#### Surface sterilization

To test the hypothesis that cyanobacteria inhabit endophytic compartments of the host mosses, some *Hylocomium* moss samples were surface sterilized. For surface sterilization, Alvarenga et al. (2024), was followed, with some modifications: 2-3 grams (about 5-10 gametophytes) of moss were manually cleaned of debris, placed into plastic 50mL centrifuge tubes, and submerged in 40mL of 0.5% Tween20 solution. Samples were ultrasonicated for a minimum of 2 minutes with a VWR ultrsonic cleaner at power 5. Liquid was removed by decanting, and replaced with 40 mL sterile de-ionized, distilled H_2_O, vortexed for 10 seconds, and discarded by decanting. Samples were then subjected twice to submersion in 1% NaOCl solution, and agitation by vortex mixer for 10 seconds at high speed. After each NaOCl application, samples were rinsed and agitated in 40 mL sterile de-ionized, distilled H_2_O. Surface sterilized samples were then again inspected for visible cyanobacterial colonization.

#### Culture of *Nostoc*-like organisms

The abundant chloroplast DNA present in host moss tissue complicates molecular laboratory and bioinformatic pipelines for characterizing cyanobacteria (Fitzpatrick et al., 2018; Song & Xie, 2020), so we attempted a cultivation-based pipeline for describing our observed *Nostoc-*like organisms during microscopic inspection. “*Nostoc*-like organisms” were defined as any filamentous bacteria that were observed in sheathed syncoenobia, especially when trichomes displayed obvious intercalary hetercytes. Media for all cultivation was nitrogen-free BG-11_0_ media (Rippka et al., 1979). Entire stems (0.5-2 cm length) of host moss were submerged into liquid media for 2-8 weeks in 15 mL Falcon centrifuge tubes at room temperatures with natural (window) lighting, followed by removal of moss tissue upon observation of phototrophic growth, to reduce further contamination. Artificial augmentation of both temperature and light levels were attempted but either did not produce additional growth or resulted in colony death. Most phototrophic growth observed in these samples were due to eukaryotic algae, but in some cases growth of *Nostoc-*like colonies was observed. These colonies of *Nostoc* or *Nostoc-*like organisms were then further separated into separate liquid media aliquots through trapping with pipette tips, retaining as little host tissue and Eukaryotic algae as possible in the process. After 1-2 months, these colonies produced additional *Nostoc* colonies of the classic, spherical morphology, which were then individually captured using micropipettes set to take up 2 microliter volumes. Colonies were then transferred to new 1.5 mL centrifuge tubes, where they were subjected to a sterile water wash and subsequent submersion in fresh BG-11_0_ media in 15 mL centrifuge tubes. These isolations were allowed to grow an additional 1-2 months, then were subject to microscopy as detailed above, and colony PCR as detailed below. As of time of writing, *Nostoc*-like organisms were successfully isolated and cultivated from 4 separate moss host species, for further DNA extraction and PCR.

#### DNA extraction

Whole-community (host and microbial) DNA was extracted using the Quick-Start Protocol DNeasy®Plant Pro Kit from QIAGEN®, following manufacturer instructions. In step 1, twice the amount of CD1 was initially added, as the specified amount of added liquid was often not sufficient to rehydrate moss tissue.

For use as additional PCR controls for whole-community nifH, DNA was also extracted from (1) liquid culture of *Anabaena variabilis* (DSM 107003), (2) leaf and stems of surface-sterilized moss *Brachythecium* sp. collected from a roadside urban grass-dominated landscape in Bayreuth, Germany, that was presumed to be abundant with nitrates and therefore low in cyanobacterial nitrogen fixation activity, and (3) surface-sterilized leaf tissue from *Solidago* sp., a common vascular plant in the region. After extraction, the DNA was checked for quality and quantity using the NanoDrop One (Thermo Scientific) and Qubit 4 assay (Thermo Fisher Scientific) and frozen at −20°C to preserve it for later testing.

#### PCR

For amplification of nifH gene, the CNF (5‘-CGTAGGTTGCGACCCTAAGGCTGA-3‘) and CNR (5‘-GCATACATCGCCATCATTTCACC-3‘) primers (Olson et al., 1998) were used as per Dıez et al. (2007). These primers are expected to amplify sequences of 359 bp length when used on cyanobacterial DNA.

NifH genes from DNA of whole-community moss+microbiome were amplified from samples of *H. splendens* collected from the University of Bayreuth Ecological Botanical Garden. Reactions were conducted in 25 μL volumes: 12.5 μL NEB Q5® High-Fidelity 2X Master Mix, 0.3 μL BSA (20 ng/μL), 1.2 μL forward and reverse primers each (10 μM), 7.3 μL PCR-grade H_2_O and 2.5 μL of template DNA. PCR conditions were as follows: initial denaturation at 98°C for 2 min, followed by 32 cycles of 98°C denaturing for 15 s, 50°C annealing for 30 s and 72°C extension for 30 s, and a final extension of 72°C for 7 min.

As isolation and growth of *Nostoc*-like colonies was extremely slow and yielded very little biomass for DNA extraction, direct colony PCR of individual *Nostoc* colonies was necessary to target and amplify full length 16S rRNA and nifH genes. Individual *Nostoc*-like spherical colonies were trapped via suction onto 20 μL pipette tips, washed in sterile water, and placed directly into 50 μL reaction volumes. PCR reagents were mixed as follows, per 50 μL reaction: 10 μL NEB Q5® 5× reaction buffer, 10 μL NEB Q5® High GC enhancer, 0.5 μL NEB Q5® HiFi polymerase, 1.0 μL mixed dNTPs (10 mM), 0.6 μL BSA (20 ng/ μL), 1.2 μL forward and reverse primers each (10 μM), 4 μL of template DNA, and PCR-grade H_2_O to 50 μL total volume. PCR conditions were as follows: initial denaturation at 98°C for 5 min, followed by 30 cycles of 98°C denaturing for 15 s, 50°C annealing for 30 s and 72°C extension for 30 s, and a final extension of 72°C for 7 min.

For 16S rRNA-targeted PCR on *Nostoc*-like colonies, protocol followed Cuscó et al., (2019), with modifications for the colony-PCR. Primers 16S-27F (5’-agrgtttgatyhtggctcag-3’) (Zeng et al. 2013) and 16S-1492R (5’-taccttgttaygactt-3’) (Klindworth et al., 2013) were used, targeting the majority of the 16s rRNA gene. PCR reagents were mixed as follows, per 50 μL reaction: 10 μL NEB Q5® 5× reaction buffer, 10 μL NEB Q5® High GC enhancer, 0.5 μL NEB Q5® HiFi polymerase, 1.0 μL mixed dNTPs (10 mM), 0.4 μL BSA (20 ng/μL), 2.5 μL forward and reverse primers each (10 μM), 4 μL of template DNA, and PCR-grade H_2_O to 50 μL total volume. PCR conditions were as follows: denaturation at 98°C for 4 min amplification, followed by 30 cycles of 98°C for 15 s, 51°C annealing for 30 s, and 72°C extension for 1 min, and a final extension of 72°C for 7 min.

PCR products were cleaned using a Nucleospin Gel and PCR cleanup column kit (Macherey-Nagel, www.mn-net.com, article-number: 740609.50), with eluted to a 14 μL volume, with final DNA concentrations between 3.8 ng/μL - 40.2 ng/μL.

#### Sequencing of nifH and 16S rRNA genes

Colony-PCR products from *Nostoc*-like cultures were then prepped according to manufacturer instructions for Nanopore-platform sequencing using either (1) the Oxford Nanopore Technologies (ONT) ligation sequencing kit V14 (SQK-LSK114) or (2) ONT Rapid sequencing V14 -Amplicon sequencing (SQK-RBK114.24), with a R10.4.1 (FLO-MIN114) generation flow cell. SQK-LSK114 kit was used in the case of 16S rRNA gene of *Hylocomium*-associated *Nostoc,* and the remaining 6 colony-PCR products were individually barcoded, multiplexed and sequenced jointly using a SQK-RBK114.24 kit. In the first sequencing run, with only *Hylocomium*-associated *Nostoc*, sequencing ran until a depth of ∼100,000 reads, with simultaneous fast-model basecalling using Guppy (MinKNOW v23.07.12, Bream v7.7.6, Guppy v7.1.4, MinKNOW Core v5.7.5). In the second, multiplexed sequencing run sequencing was run until total depth of ∼38,000 reads, without basecalling, which was conducted separately with Dorado basecalling software (MinKNOW v25.05.12, Bream v8.5.4, Dorado v7.9.8, MinKNOW Core v6.5.13). In both cases default minimum Q-scores of 8 were used to define passing reads.

### Bioinformatics

#### Processing raw reads to consensus sequences

Demultiplexing and trimming of adapters and barcodes of Nanopore reads from colony-PCRs of *Nostoc*-like cultures was done using the Dorado basecalling softare. Resulting reads were then re-oriented 5’-3’ where necessary, using VSEARCH (Rognes et al., 2016) v2.30.0, against the Silva 138.2 (Quast et al., 2013) Genus-level training dataset, formatted for DADA2 software (“silva_nr99_v138.2_toGenus_trainset.fa”, available at www.arb-silva.de/current-release/DADA2/1.36.0/SSU). Re-orientation of nifH reads was conducted against the custom nifH database of nifH sequences from Kubota et al. (2023).

Resulting reads from nanopore sequencing were typically numerous and very noisy, aligning poorly due to numerous erroneous insertions, paralogs, and co-cultured microorganisms. In some cases, BLAST(Basic Local Alignment Search Tool)-family algorithms (Altschul et al., 1990) have been found to perform effectively for aligning reads with common error patterns of long-read metabarcoding data (Tedersoo et al., 2021, 2022). Therefore, as a first filtering step before full alignments, we used the BLAST-suite tools, specifically blastn and blastx programs to find high quality matches to *Nostoc* 16S sequences from the Silva 138.2 database (see link above), and to a custom database of nifH amino acid sequences. Following Hu, Wang, et al. (2024), our custom database of nifH sequences was a concatenation of NifH sequences from (1) 963 culture-derived genomes (Koirala and Brözel, 2021), the Greening lab NifH database (https://doi.org/10.26180/c.5230745) and the Nathoo database (Suquilanda-Pesántez et al., 2022). Following this filtering step, reads were aligned de novo using MUSCLE 5.1 with default settings (Edgar, 2022). Vertical gaps were removed using CIAlign (Tumescheit et al., 2022), using a insertion minimum percentage of 70%, of all gap sizes (insertion minimum size > 0, insertion minimum flank size > 0).

Both 16S (Sun et al., 2013) and nifH (Thiel, 1993; Thiel et al., 1995) can be present in multiple unique copies within a single organism. To detect this possible paralogous, intra-genomic variation, trees of full alignments of amplicon sequences were constructed using FastTree 2.2 with default settings (Price et al., 2010), and visualized with ARB 7.0 (Ludwig et al., 2004). Unrooted trees were searched for obviously separate clades (example shown in Supp. Fig. 1). If present, these reads from these subtrees were again aligned individually using MUSCLE, and manually curated for removal of poorly aligned sequences and ends. If obviously separate clades were not present in the phylogenetic tree, the full alignment of 16s or nifH was assumed to represent a single representative gene sequence for that organism. In either case, consensus sequences were generated using Hidden Markov models (HMM) from each (sub)alignment, using the hmmbuild and hmmemit commands, from the HMMER v3.3 package (Eddy, 2011), using simple majority-rule decision for consensus-sequence bp calls. Since nifH Nanopore sequences were not preliminarily taxonomically filtered to *Nostoc*, unlike 16S (see above), resulting nifH consensus nucleotide sequences were at this point searched against the full NCBI Core nucleotide BLAST database (<u>blast.ncbi.nlm.nih.gov</u>), for highly similar matches using the online blastn graphical user interface, to confirm likely Nostocaceae origin.

#### NifH phylogeny

Our nifH phylogenetic analysis was intended to follow - and build upon - Ininbergs et al. (2011) and Kubota et al. (2023). All pertinent nifH sequences from both studies were provided by authors (Masayuki Kubota, pers. comm.), combined with nifH consensus sequences from our cultures. Alignment was conducted using Muscle v5.1 with default settings, and tree was constructed using FastTree2 using default settings.

#### 16S Nostocaceae phylogeny

For taxonomic identification of full-length 16S consensus sequences from cultured *Nostoc*-like organisms, a de novo phylenogenetic tree of full-length 16S sequences for Nostocaceae was constructed. The full Cyanoseq (Lefler et al., 2023) v1.2 phylogenetic tree (https://zenodo.org/records/7864137) was visualized in ARB v7.0, and sequences of all members of Nostocaceae were selected, including sequences from closely allied clades Tolypotrichaceae, Aphanizomenaceae, and Nodulariaceae (Strunecký et al., 2023).

Additionally, 708 sequences of full or near-full length 16S from NCBI genbank that were identified as high-quality *Nostoc* sequences by Cyanoseq v1.3 were provided by the authors for use in our 16S tree (Forrest Lefler, pers. comm; full cyanobacterial dataset available at https://zenodo.org/records/13910424), along with several of 16S reference sequences of interest: *Nostoc azollae* (NC_014248.1) (Ran et al., 2010), *Nostoc calcicola* SAG 1453-1 (KM019926) and *Nostoc insulare* SAG 54.79 (KM019927). Full 16S sequence of *N. azolla*e was not available as a stand-alone NCBI nucleotide accession, so 16S copies were extracted from reference genome with barrnap (https://github.com/tseemann/barrnap). *Scytonema pachmarhiense* (MH260366) was added as an outgroup for rooting of phylogenetic trees. In total, 856 sequences were included in our Nostocaceae tree. Reads were aligned using SSU-align v0.1.1 (Nawrocki, 2009), using default Bacterial models. Tree was constructed using FastTree2. Full alignment FASTA and tree Newick files are available in supplementary materials. Tree topology was visualized with ARB 7.0, and final cosmetic additions to tree labels and highlighting of taxa of interest were done using Inkscape v1.3.2 for GNU linux.

## Results

### Microscopy

*Nostoc*-like organisms were observed in 5 moss species, from 4 sites (Figures 2-6): *Orthotrichum lyelli* from Northern Fraconianjura bei Neustädtlein*, Hylocomium splendens* from Waldstein*, Zygodon rupestris* and *Pseudoamblystegium subtilis* from Pfaffenberg bei Mettendorf, and *Homalia trichomanoides* from Schwarzachklamm bei Gsteinach. Within each moss, position of *Nostoc-*like colonies were variable but were most common in the leaflet axils of the moss gametophytes. Most notably in *Orthotrichum lyelli,* the position of *Nostoc–*like colonies were more variable, occurring in the leaf axils but also often on leaf laminas.

**Figure 2.**
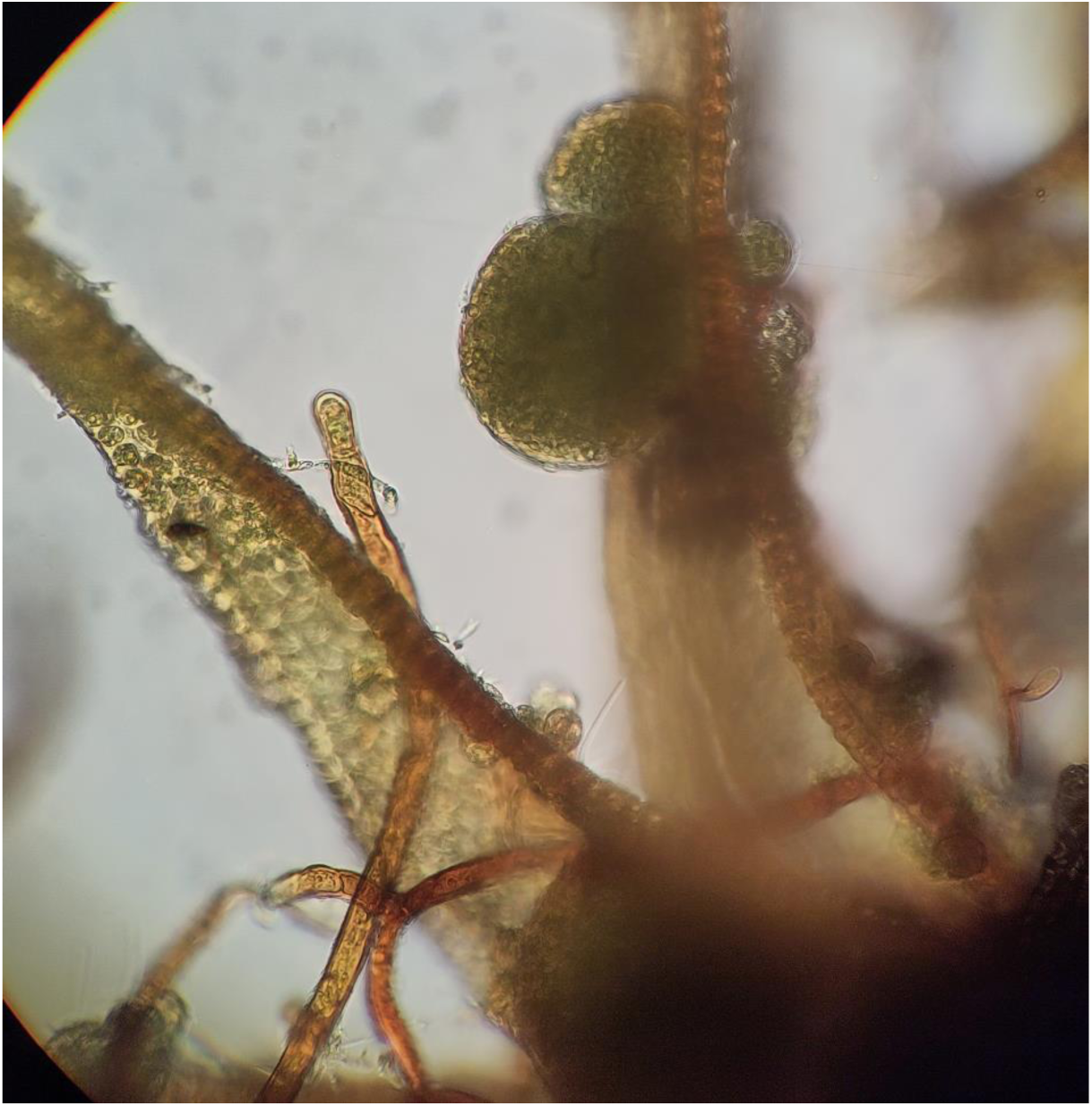
Colonies of *Nostoc-*like organism observed from *Orthotrichum lyelli* at 400× magnification.

### Surface sterilization

Surface sterilization of *Hylocomium splendens* fronds greatly reduced the amount of visible colonization by cyanobacteria and other phototrophs (Fig. 7). Generally, surface sterilized leaf surfaces were cleared of visible colonies, though occasionally some microbial cells were observed (Fig. 7c). *H. splendens* stems still appeared to consistently harbor extensive significant microbial biomass following surface sterilization, possibly cyanobacteria because of their fluorescence patterns (Fig. 7d).

### Culture of *Nostoc*-like organisms

Out of 5 observed *Nostoc*-like organisms, 4 strains were successfully isolated. Successful *Nostoc-*like isolates were from: *Hylocomium splendens* (Fig. 3)*, Homalia trichomanoides* (Fig. 5)*, Pseudoamblystegium subtile* (Fig. 6)*, and Zygodon* sp. (Fig. 4). The *Nostoc-*like organism from *Orthotrichum* moss (Fig. 2) failed to show any growth and typically became chlorotic, despite abundant growth on the collected moss tissue. The *Pseudoamblystegium-*associated *Nostoc-*like organism was lost shortly after transfer into liquid culture, and was overtaken by an unidentified, co-cultured non-filamentous cyanobacterium. This may have resulted in a failed PCR amplification of nifH, though some 16s data from the *Nostoc*-like organism were recovered (see below).

**Figure 3.**
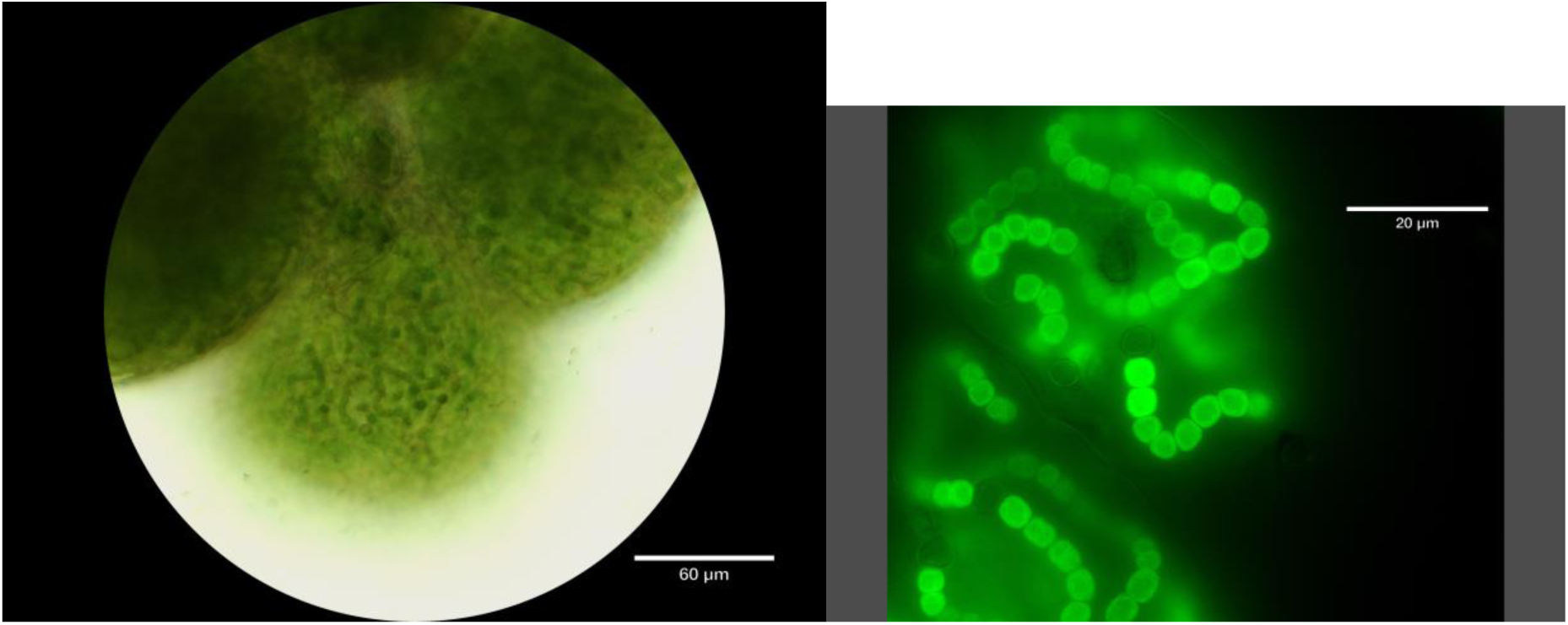
Colonies of *Nostoc-*like organism after isolation from *Hylocomium splendens.* (a) light microscopy, at 400× magnification and (b) 1000× magnification using epifluorescence with Cy5 filter setup (see methods), showing trichomes and heterocysts.

**Figure 4.**
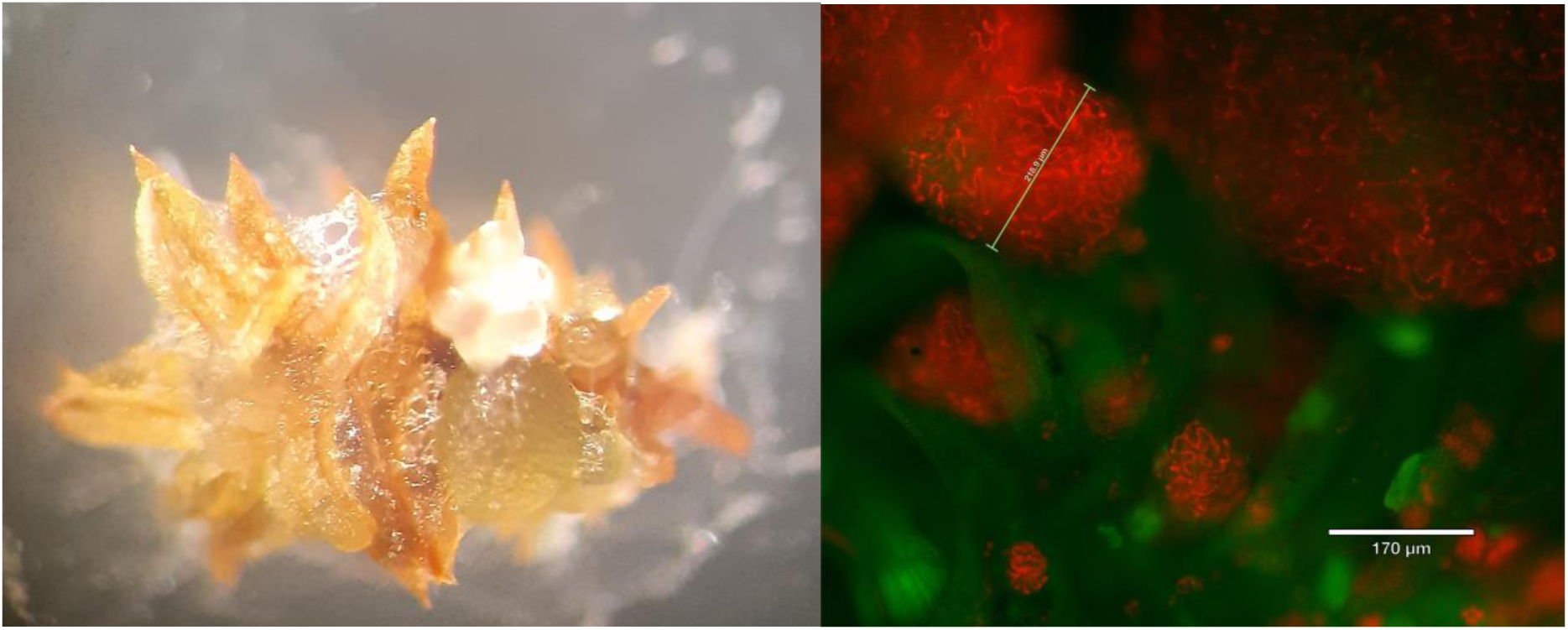
Colonies of *Nostoc-*like organism observed from *Zygodon rupestris* (a) at 50× magnification with binocular dissecting microscope show heavy colonization of moss caulid. (b) at 400×, same colonies using using epifluorescence with Cy5 filter setup for detection of cyanobacteria, overlaid onto FITC setup for background moss tissue (see methods).

**Figure 5.**
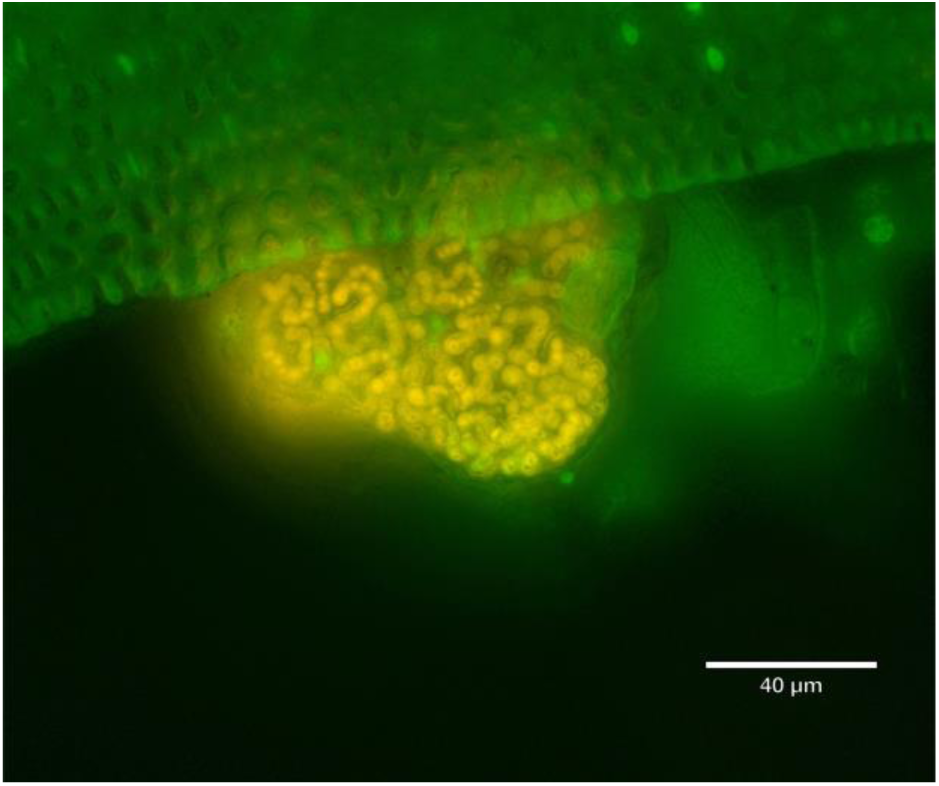
Colonies of *Nostoc-*like organism observed from *Homalia trichomanoides at* 400× using epifluorescence with TXRED filter setup for detection of cyanobacteria, overlaid onto FITC setup for background moss tissue (see methods).

**Figure 6.**
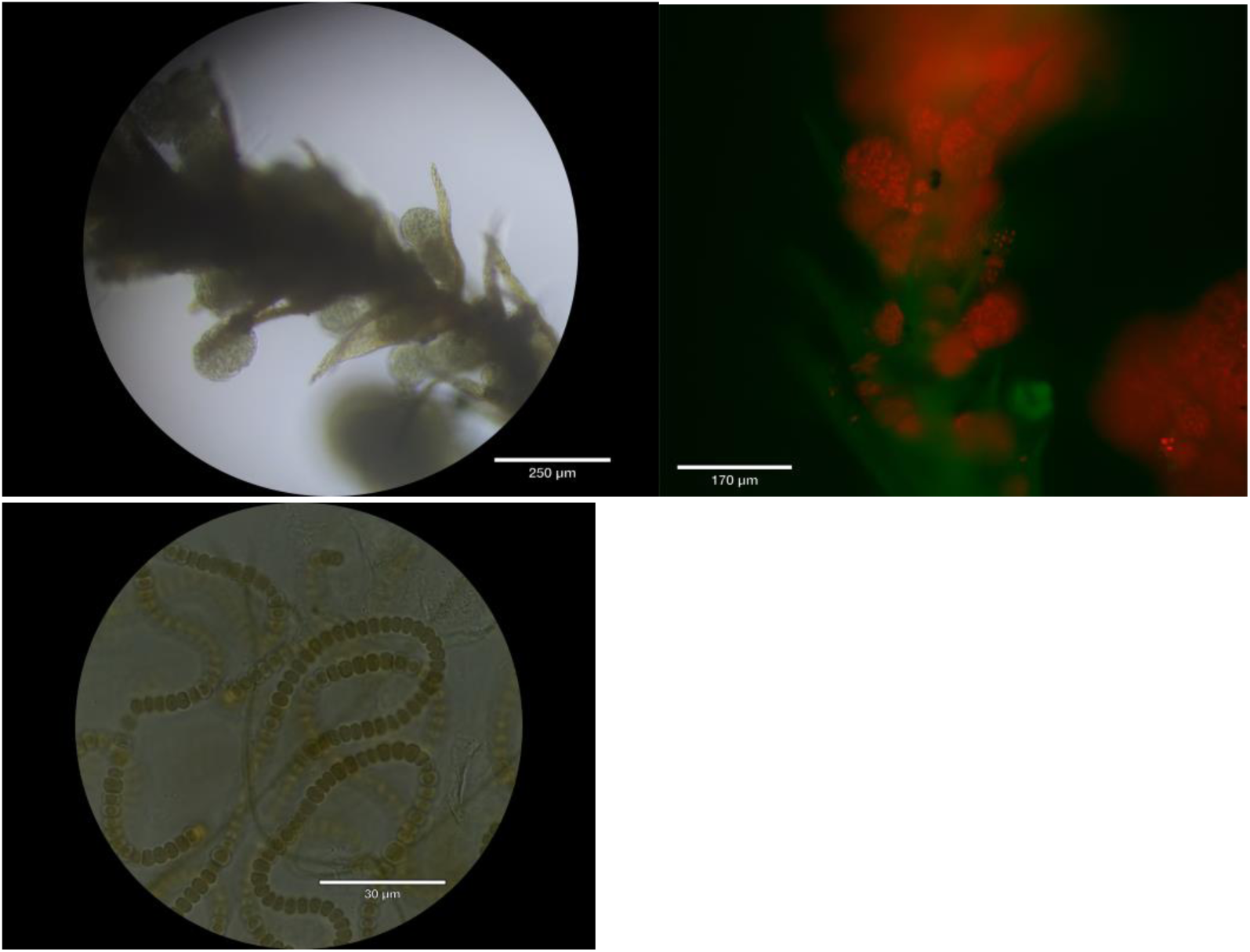
Colonies of *Nostoc-*like organism observed from *Pseudoamblystegium subtile.* (a,b) abundant colonization by nostoc colones on caulid at 100× magnification with light and using epifluorescence with Cy5 filter setup for detection of cyanobacteria, overlaid onto FITC setup for background moss tissue (see methods). (c) 1000× magnification of same organism, showing trichomes and heterocysts.

**Figure 7.**
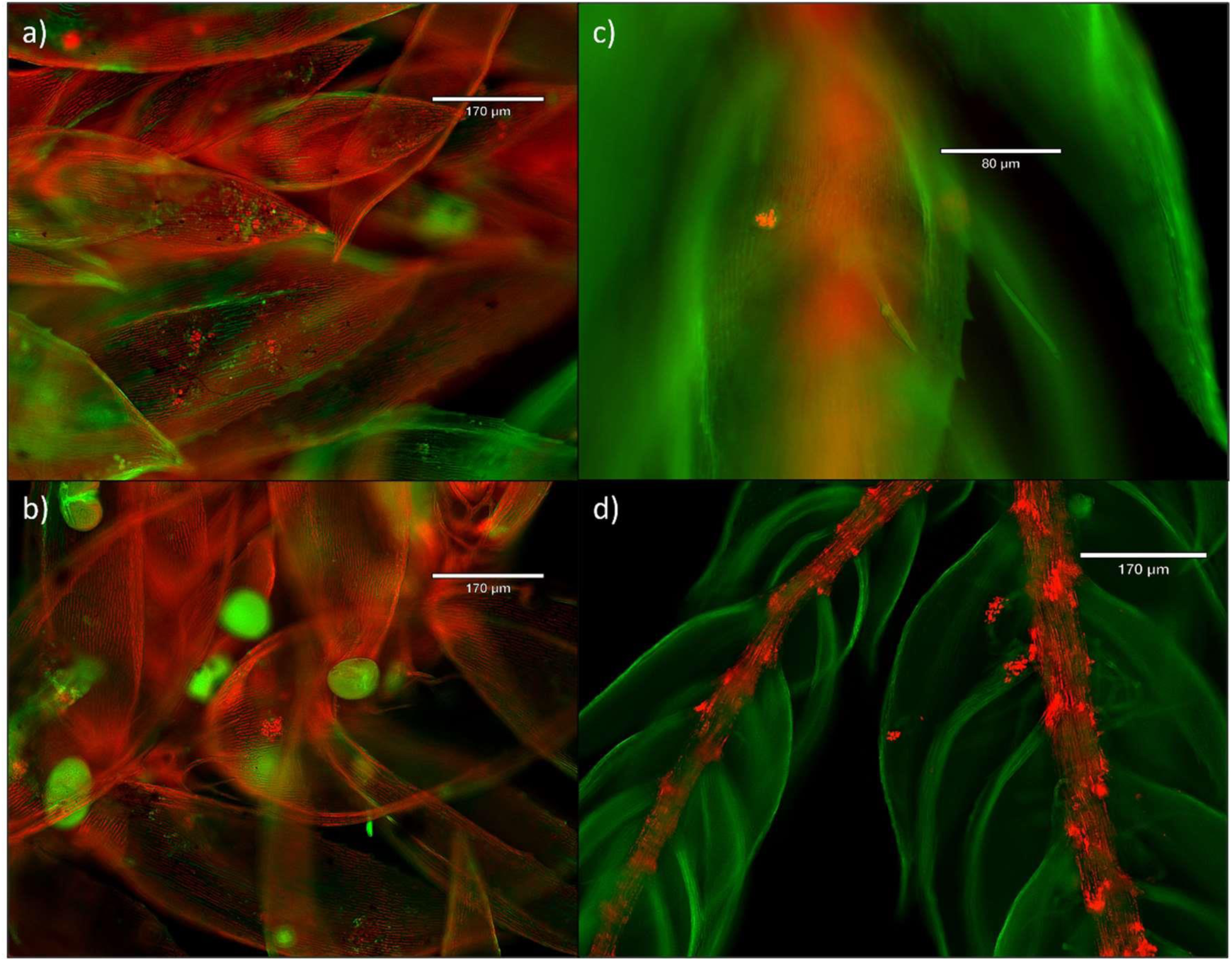
Microbiota of leaves and stems of *Hylocomium splendens,* using epifluorescence with Cy5 filter setup for detection of cyanobacteria, overlaid onto FITC setup for background moss tissue (see methods). (a-b) fluorescence of cyanobacterial and eukaryotic algae on untreated moss leaves. (c,d) after surface sterilization, eukaryotic-algal and cyanobacterial cells on leaves are mostly no longer visible, but colonies on stems remain visible, possibly indicating endophytic lifestyle within stems.

### Whole-community amplification of nifH gene

Culture-free amplification of cyanobacterial nifH from *Hylocomium* moss tissue yielded two size classes of non-chimeric amplicons, one at the expected size of slightly less than 400 bp, and anot her with a size of >500 bp. (Fig. 8). Based on PCR controls from *Nostoc-*free plant tissues and from our *Anabaena* culture, we presume that the smaller size class represents the intended target, cyanobacterial nifH, and the other represents non-target homologous genes from host moss tissue (Angel et al., 2018). Both amplicon types were observed in all portions of *Hylocomium* moss tissue (low, middle, upper), indicating possible cyanobacterial colonization of moss tissues even when not visible through standard microscopic inspection.

**Figure 8.**
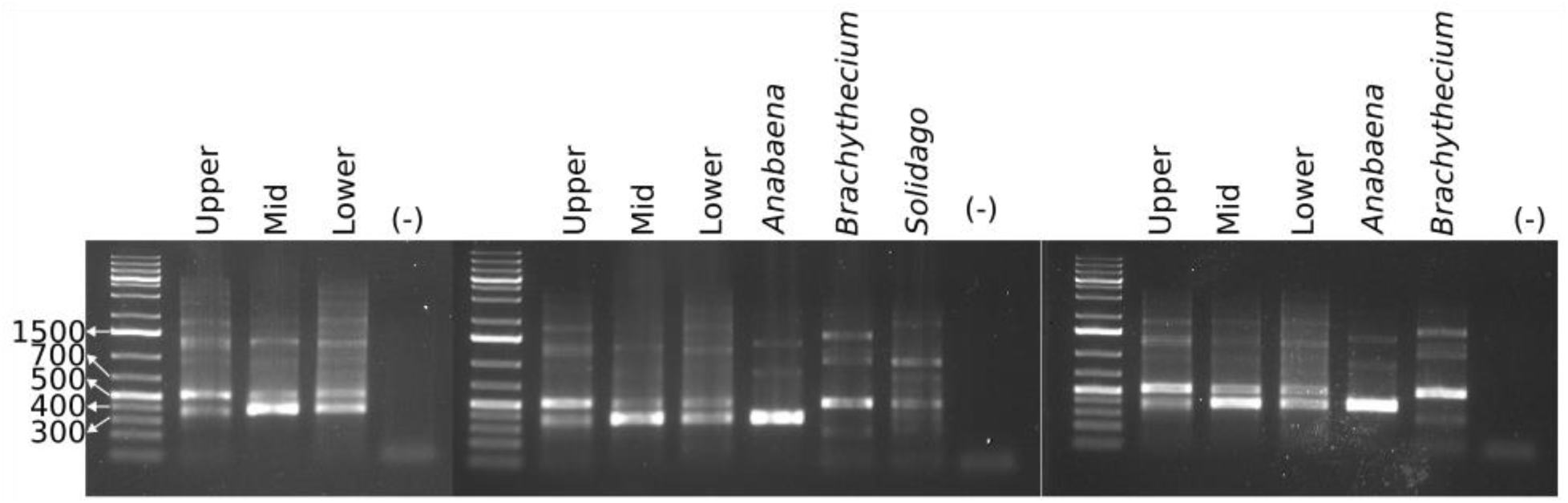
Electrophoresis gel ol PCR of nifH region, using cyanobacterial specific nifH primers on whole community DNA extracted from *Hylocomium splendens*. Bands above 700 bp in size are presumed to be chimeric in origin due to overamplification. Bands in the region of 500 bp are hypothesized to be embryophyta homologs of nifH, due to their presence in surface-sterilized, urban *Brachythecium* moss and surface-sterilized *Solidago* leaves, and their absence in axenic *Anabaena* culture. Amplicons of size ∼400 are presumed to be the intended the target of PCR reaction, cyanobacterial nifH, indicating the possible presence of nitrogen-fixing cyanobacteria in these moss tissues.

### Phylogenetic placement of nifH genes from cultured *Nostoc*-like organisms

We detected nifH sequences from 3 of the 4 *Nostoc*-like organisms: *Hylocomium-, Zygodon-,* and *Homalia-*associated *Nostoc-*like organisms. Despite differentiated cells resembling heterocysts (Fig. 6), PCR-amplification of nifH from our culture of *Pseudoamblystegium-*associated *Nostoc-*like organism did not yield any sequence-able product, probably due to overgrowth by a non-nitrogen fixing cyanobacteria (see above). The topology of our nifH tree recaptured the major clades of Kubota et al. (2023) and Ininbergs et al. (2011) with some minor differences (Fig. 9, supp. Fig. 2). The most important distinction was the inclusion of the *Stigonema* nifH cluster within Nostoc cluster I. In *Zygodon-* and *Homalia-*associated cultures each appeared to host 3 alternative nitrogenases, each falling into a separate clade on the Kubota-Ininbergs nifH tree. Far fewer high-quality nifH sequences were recovered from the culture of our *Hylocomium-*associated *Nostoc*-like organism,in comparision to our two other *Nostoc-*like cultures (supp. Fig 1), but these were sufficient to generate a single candidate consensus sequence for nifH in this organism.

**Figure 9.**
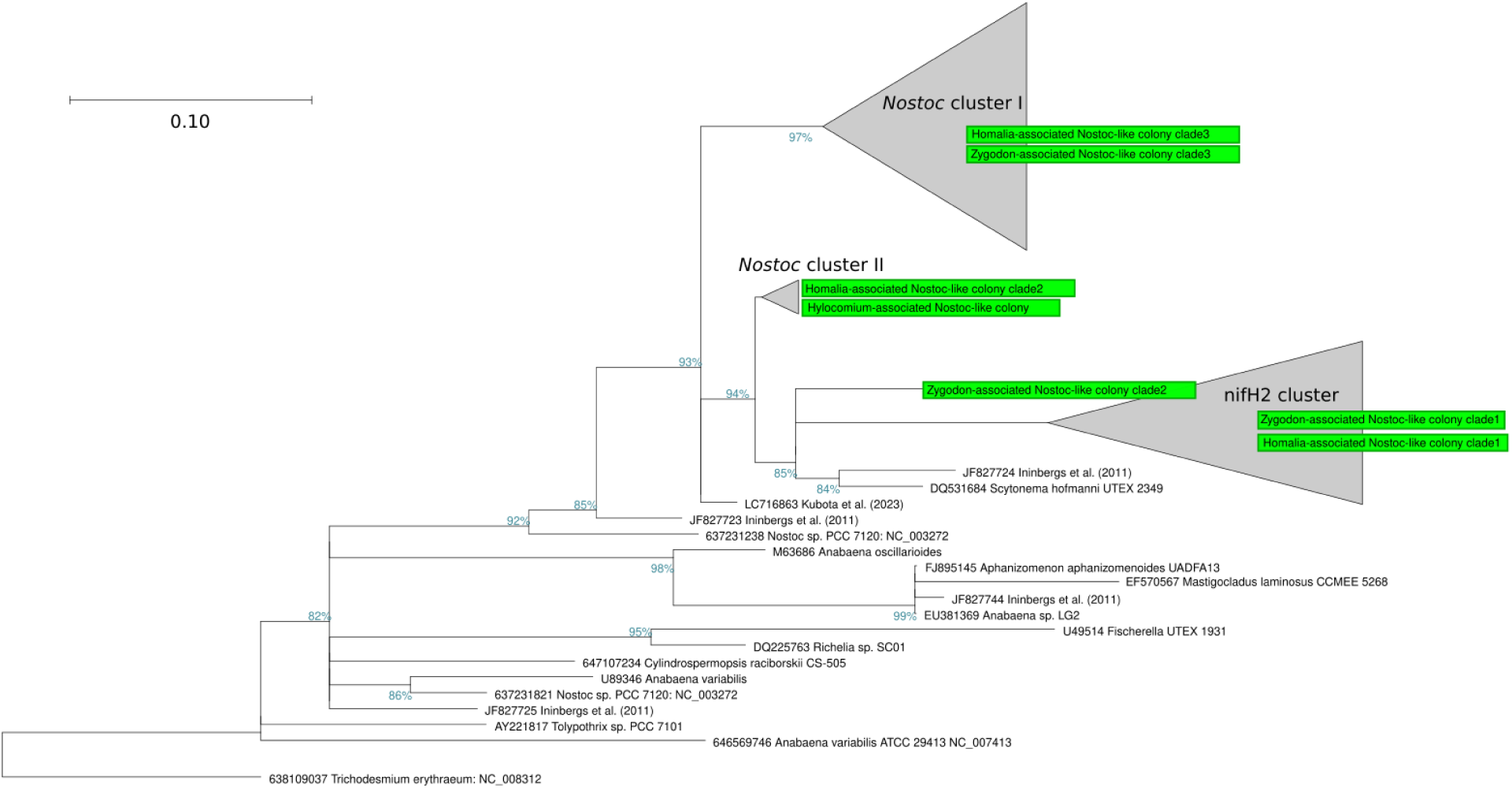
nifH gene phylogeny of select members of Nostoceae. Tree is constructed from de novo alignment of nifH sequences detected in this study from *Nostoc-*like organisms (green) and reference sequences from Ininbergs et al. (2011) and Kubota et al. (2023). *Zygodon*- and *Homalia*-associated *Nostoc-*like organisms both hosted multiple distinct copies of nifH. Full-size tree with all clades expanded, including placement of “Stigonema Cluster”, is available as Supp. Fig. 2.

### Phylogenetic placement of 16S genes from cultured Nostoc-like organisms

Full-length 16S consensus sequences were determined for all 4 successfully cultured *Nostoc*-like organisms (Fig. 10, green circles). Only one consensus Nostocaceae 16S sequence was detected for each of the cultures. In the case of *Pseudoamblystegium,* 16S sequence data were overwhelmingly non-*Nostocaceae* cyanobacterial reads that were not identifiable beyond phylum-level (Cyanobacteriota), likely due to co-cultured cyanobacteria (see above). However some high-quality 16S reads remained for identification of the *Nostoc*-like organism originally observed for this sample.

**Figure 10.**
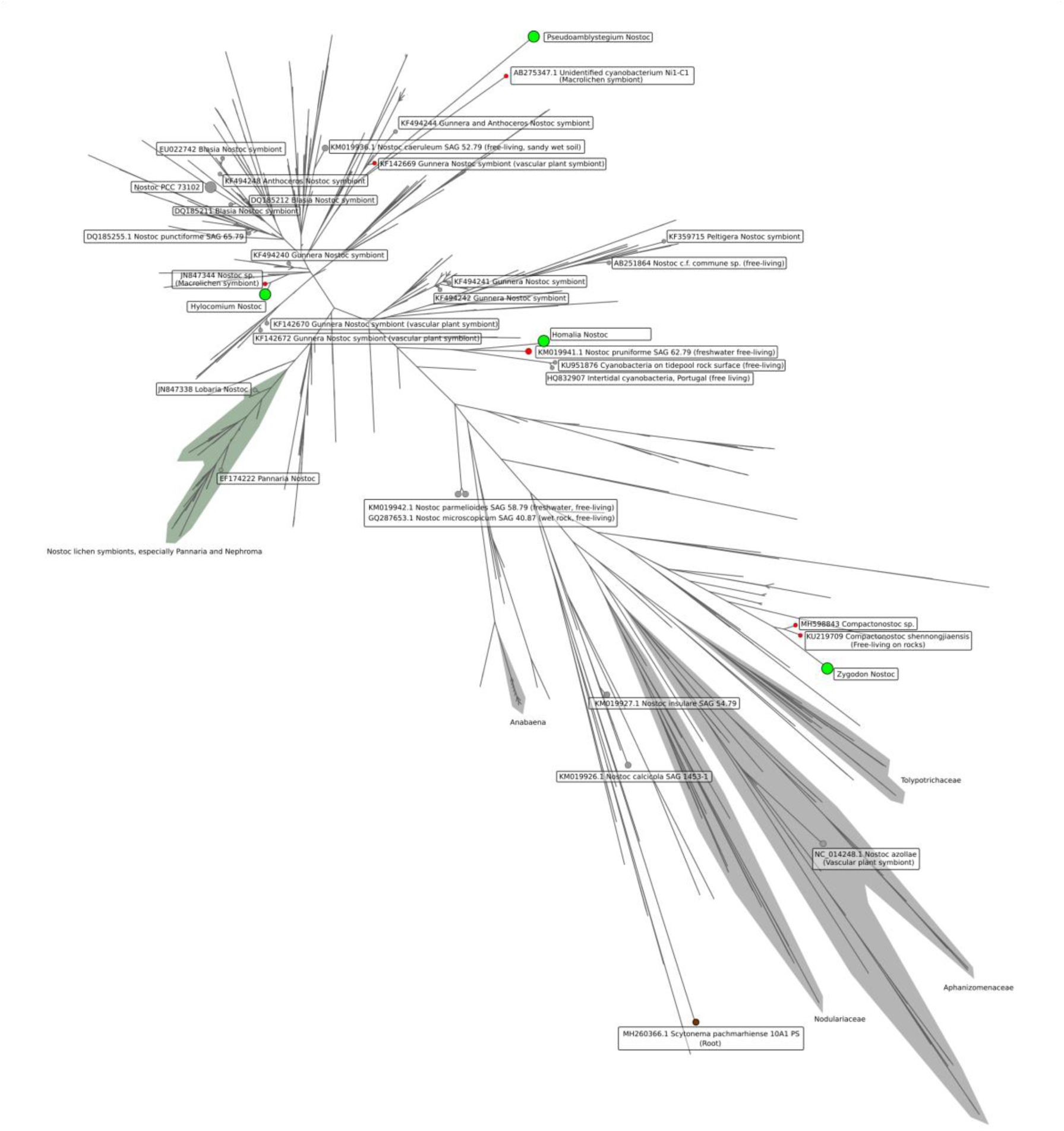
De novo 16s Nostaceae phylogeny constructed from 856 full length 16S sequences from the Cyanoseq database (see methods), selected additional reference sequences, and four 16S consensus sequences from moss-associated *Nostoc*-like organisms from the present study (green circles). Nearest neighbors to moss-associated *Nostoc*-like organisms are shown by smaller red circles, and selected additional reference sequences are labeled by gray circles.

Observed *Nostoc*-like moss symbionts were placed throughout Nostocaceae (Fig. 10, Table 2), though nearest neighbors were in several cases previously known symbionts: in *Pseudoamblystegium* and *Hylocomium* mosses, the moss-associated *Nostoc* were closely related to endosymbionts of macrolichen-symbionts (Miura & Yokota, 2006; Olsson et al., 2012) and the vascular plant *Gunnera magellanica* (Fernández-Martínez et al., 2013). For *Zygodon* moss, the cyanobacterium we observed was most closely related to the recently described genus *Compactonostoc* (Cai et al., 2019). *C. shennongjiensis* was reported as free living on rocks, another secondary habitat known for *Zygodon* species. For the moss *Homalia trichomanes*, the closest relative was the freshwater-dwelling *Nostoc pruniforme,* as per Experimental Phycology and Culture Collection of Algae (SAG) strain 62.79, and other free-living *Nostoc* strains. Closest matches to other 16S sequences from Nostocaceae are shown in Table 2.

**Table 2.** Closest sister taxa to observed *Nostoc-*like organisms observed from mosses in this study, based on our de novo 16S tree of Nostocaceae (Fig. 10). When a very close match is available, only one taxa is listed. When a close match is not obvious, the two nearest neighbor taxa are listed.

| Moss host | Sister taxa / nearest neighbor |
| --- | --- |
| *Hylocomium splendens* | JN847344 *Leptogium*-associated *Nostoc* sp. (macrolichen symbiont) |
| *Homalia trichomanoides* | KM019941 *Nostoc pruniforme* SAG 62.79 (freshwater, free-living) |
| *Pseudoamblystegium subtile* | 1. KF142669 *Gunnera*-associated *Nostoc* symbiont (vascular plant symbiont)<br>2. AB275347.1 Unidentified cyanobacterium Ni1-C1 (unidentified-macrolichen symbiont) |
| *Zygodon dentatus* | 1. MH598843 *Nostoc* sp., unpublished, classified as likely *Compactonostoc* by Cyanoseq<br>2. KU219709 *Compactonostoc shennongjiaensis* (Free-living on rocks) |

## Discussion

Here we have documented additional instances symbioses of *Nostoc* and *Nostoc-*like organisms with several mosses from the region of middle and upper Franconia, Germany, utilizing a microscopy-, culture-, and barcoding-based approach. Our survey therefore expands the known range of these symbioses further into southeastern Germany. We have also expanded the range of known moss hosts for *Nostoc* to include two non-feather mosses, in the family Orthotricaceae (order Orthotrichales), represented here by *Zygodon* and *Orthotrichum.* Additional new host moss families within *Hypnales* were also documented: *Pseudoamblystegium subtile* (Amblystegiaceae) and *Homalia trichomanes (*Neckeraceae).

The finding of Nostoc symbiosis in forest-floor *Hylocomium splendens* deeper into central Europe was predicted, since the well-known host moss species *H. splendens* and *Pleurozium schreberi* are very common in the region. Nevertheless, the extension of the geographical range of this well-known symbiosis further emphasizes their likely importance to montane forest ecosystems. Given the documented importance of moss-hosted cyanobacterial nitrogen in other systems (discussed above), the contribution of these symbioses to the primary productivity of montane forest ecosystems of our region of Central Europe is also likely to be significant. This argues for the protection of forest-floor moss layers and canopy mosses in the montane forests of the region as part of maintenance of these forest stands.

There are obvious disadvantages to a culture-based approach, primarily that presumably many *Nostoc*-like organisms were not successfully cultured. Cultures that were successful were extremely slow-growing, and were in one case not mono-algal. Generally, target *Nostoc-*like organisms were often outcompeted in culture by other photoautotrophs. *Nostoc*-like organisms exist in many other life stages besides colony (syncoebium) stages (Mollenhauer, 1986), including single-cell and single-trichome life stages. *Nostoc* present on mosses in these life stages probably often escape detection by standard light microscopy. Our PCR-tests of moss tissue with cyanobacterial nifH genes (Fig. 8) support this, indicating that cyanobacteria are likely present in most aerial tissues of *Hylocomium* even when not detectable with microscopy. As such, the organisms cultured here likely greatly under-represents the diversity of *Nostoc*-like organisms in mosses in our region.

Despite the limitations of our culture- and microscopy-based approach, we documented new moss hosts for *Nostoc*-like organisms. Particularly of interest are host species *Zygodon dentatus* and *Orthotrichum lyellii,* both of Orthotricaceae and both of which fall outside of the category of feather mosses (order Hypnales). Feather mosses and *Sphagnum* are traditionally understood to be the hosts for moss-*Nostoc* symbioses, but we show here that such symbioses may be occurring in other distantly-related moss groups.

Importantly, these two new host mosses (*Zygodon* sp. and *Orthotrichum lyellii*) were arboreal mosses. Four out of five moss hosts in which we observed to host *Nostoc-*like organisms were epiphytic on tree bark (Table 2). We did not conduct systematic sampling for *Nostoc*-like organisms, so we cannot quantify the relative abundance of the symbiosis in canopy-dwelling mosses in our region vs. forest floor mosses. However, this is supportive of the hypothesized importance of this symbiosis in forest canopies (Lindo & Whiteley, 2011), and we hypothesize that N-fixing cyanobacterial symbionts may provide an important mechanism for allowing moss communities to thrive and compete in the very challenging microclimate of bark. Cyanobacteria in arboreal mosses have been observed previously, i.e. in the coastal temperate rain forest of North America by Lindo and Whiteley (2011) and in a subtropical montane cloud forest in southwestern China for the species *Homaliodendron montagneanum*, *H. scalpellifolium* and *Thuidium cymbifolium* (Fan et al., 2022). Physically smaller and more ephemeral mosses such as *Zygodon* spp. or *Pseudoambylstegium* spp., as well as initial or young stages of larger mosses like *Orthotrichum,* may especially benefit from this symbiosis, as smaller mosses may not be able to access biologically available nitrogen from atmospheric deposition or stemflow as well as larger mat-forming mosses. If this is true, we would predict symbiosis with cyanobacteria to be especially prevalent on the smaller species as hosts, as we observed for *Zygodon, Pseudoamblystegium subtile* and *Homalia*, all of which are comparatively physically small mosses of their families. Whereas the genus *Orthotrichum* is widespread and common in our region, the genus *Zygodon* is rare in central Europe and occurs ephemerally in very scattered mats at sites with a high humidity. *Zygodon* has comparatively small gametophytes and may be at a competitive disadvantage compared to larger, co-occuring species of *Orthotrichum*, and the moss-*Nostoc* symbioses may be especially important for these mosses for continued survival. Additionally, *Zygodon* spp. and *Orthotrichum lyellii* produce large amounts of multicellular gemmae for dispersal. This continual production of gemmae may create additional demand on these mosses for organic nitrogen, and therefore symbioses as we observe here.

We found no clear phylogenetic signal among the symbiotic *Nostoc*-organisms for which the 16S rRNA gene were successfully sequenced - symbionts were placed throughout the Nostocaceae family (Fig. 10, Table 2). Nearest matches to other *Nostoc* full length 16S sequences were from various ecological niches, both symbiotic and free-living. Additionally, according to 16S sequence data, our *Nostoc-*like organism isolated from *Zygodon* is most closely related to the recently-described genus *Compactonostoc* (Cai et al., 2019). This genus of *Nostoc-*like organisms has to-date been observed from freshwater free-living states (Cai et al., 2019; Caro-Borrero et al., 2024), but our observation implies that members of *Compactonostoc* may be capable of plant-symbiosis.

Another benefit of our culture-based approach is that we are able to link taxonomic identity (full-length 16S sequences) to a functional gene (nifH) within single organisms. This has enabled the observation of possible alternate nitrogenases (3 distinct nifH copies) in the genomes of 2 of our moss-associated *Nostoc-*like organisms. Alternative nitrogenases have been previously observed from members of Nostocaceae (Thiel, 1993; Thiel et al., 1995), and are thought to perform supplementary or complementary roles, allowing diazotrophy in different oxygen levels, or the use of alternative metal co-factors (vanadium or iron-only) in place of molybdenum (Zehr et al., 2003). It is worth noting that our only terrestrial moss (*Hylocomium splendens*) did not appear to host *Nostoc* with alternate nitrogenases, only our arboreal *Nostoc* were observed to have alternate *NifH* versions. We speculate that in the forest canopy environment, the availability of molybdenum may be limiting, requiring alternate nitrogenases with other co-factors. Alternatively, the dynamic microclimate of the forest canopy may select for homologs of nitrogenase genes with different environmental optima. Either possibility would enable diazotrophic cyanobacteria to fix nitrogen and continue growth under a wider range of conditions.

The possibility that moss-associated cyanobacteria host multiple nitrogenases has implications for the study of moss-associated *Nostoc* and their ecological services. For example, Warshan et al. (2016) and Arróniz-Crespo (2022) attribute relative importance in terms of biological nitrogen fixation to different cyanobacterial groups in boreal forests, based on differential abundances of sequences falling into nifH clades defined by Ininbergs et al. (2011). However, we show here that great care should be taken when assigning taxonomy using these nifH clusters. Warshan et al. (2016) and Arróniz-Crespo (2022) suggest *Stigonema* as the most important group contributing to fixation of nitrogen in boreal forest systems based on high relative abundance or high levels of expression of nifH genes aligned to the “Stigonema cluster” of nifH, sensu Ininbergs et al. (2011). Our results indicate that the *Nostoc* strain cultured from *Homalia trichomanoides*, closely related to *Nostoc pruniforme* SAG 62.79 (Fig. 10), has a version of nifH that falls within this “Stigonema” cluster, but also hosts two other nifH genes from other clusters sensu Ininbergs et al. (2011) (Fig. 9 and Supp. Fig. 2).

We also observed the possible endophytic colonization of moss stems by cyanobacteria (Fig. 7). Among mosses, cyanobacterial endophytes are only previously known in *Sphagnum,* which hosts *Nostoc* and other bacteria in specialized structures known as hyalocytes, or dead, hyaline leaf cells (Carrell et al., 2022). Moss leaflet lamina are typically only one cell thick (Roberts et al., 2012), probably precluding abundant endophytic colonization of leaves by microbes that is commonly observed in vascular plants (Bodenhausen et al., 2013; Rodriguez et al., 2009).

Stems (‘caulids’) and rhizoids of mosses are multicellular, and are therefore presumably the site of colonization of much of the extensively documented, diverse non-cyanobacterial endophytes observed in mosses (Tveit et al., 2020; U’Ren et al., 2010). If the observed cells are indeed cyanobacteria occupying the stem tissue of mosses, it points to an abundant symbiosis. We found epiphytic syncoebia relatively rarely in our mosses, so the possibility exists that much of the relatively high rates of nitrogen fixation by cyanobacteria among mosses in our region as noted by Groß et al. (2024) may be achieved by these potential endophytic symbionts instead. Endophytic colonization by *Nostoc* and other cyanobacteria is well documented in other plant-and algal-symbiotic systems, including in hornwort and liverwort hosts (reviewed by Adams and Duggan (2008)). Further experimental work is needed to confirm the cyanobacterial nature of these cells, and if they are capable of nitrogen fixation.

## Conclusion

Here we extend the known geographical range of the classical feather moss *Hylocomium*/*Nostoc* symbiosis further into the region of central Europe, highlighting the importance of moss layers for nitrogen cycling in montane forests of central Europe. We expand the known host range of to include additional non-feather mosses (*Orthotrichum* and *Zygodon*), as well as to other feather mosses not previously observed to host *Nostoc* symbionts (*Ambylestegium, Homalium*). We note that multiple paralogs of nifH can be found within some *Nostoc-*like organisms, which may denote adaptation to the dynamic environment of the forest canopy, but which also implies that great care be applied when attempting to identify these Nostocaceae symbionts with nifH gene sequence data. We also report a potential site of high cyanbacterial colonization, namely the endophytic compartment of the “stem” (caulid) of mosses. If occurring, this implies an abundant, physically-protected symbiosis but not with *Nostoc,* or at least not in the large-colony stages.

Taken together, these results help to bridge the gap between two commonly observed forest moss symbioses: (1) few-to-few, feather-moss/*Nostoc* symbioses, and (2) many-to-many moss/cyanobacterial-community symbioses. From our survey, the former appear to occur in our region but relatively rarely. Based on other reports (cited above), the latter class of broader cyanobacterial symbioses is apparently unbiquitous in mosses where nitrogen is limiting, including in our region. The question now arises as to whether there are particular advantages for mosses to entering into symbiosis with *Nostoc-*like organisms in particular, or whether these feather-moss/*Nostoc* symbioses are simply another case of very common moss-cyanobacterial symbioses. It is also very possible that complex and diverse cyanobacterial communities in mosses have been understudied historically by researchers when highly visible and charismatic *Nostoc* colonies are present. However, in the course of our surveys we noted *Nostoc-*like colonies more often in arboreal habitat, and in locally uncommon mosses that were relatively specialized tree-epiphytic mosses such as *Zygodon*. As such we suggest that host-symbiont specialization could be occuring in the special habitat of the forest canopy. However, quantitative surveys to compare the prevalence of *Nostoc-*like organisms between the among habitats are still mostly lacking, and the unique benefits of such specialized canopy symbioses, if any, are yet to be explored.

## Declarations

### Ethics approval

This article does not contain any studies with human participants or animals performed by any of the authors.

### Consent to participate

not applicable

### Consent for publication

not applicable

### Availability of data and material

code and supporting files are available at the associated github.com repo in the folder “suppData”, at the following link: <https://github.com/danchurch/scalingUpByLookingCloser_Feulner_etal/tree/main/suppData>. Demultiplexed, raw Nanopore reads are available at NCBI SRA, from the bioproject PRJNA1401537.

## Author Contributions

MF conceived of the survey topic and design. All authors contributed to field work. MF, US, FJPD, and DCT contributed to microscopic imaging and culturing of microorganisms. JM contributed the hand-drawn illustration of Hylocomium moss and Nostoc (figure 1). MF and US identified moss species. JM, FJPD and DCT conducted laboratory experiments. DCT conducted DNA sequencing and wrote bioinformatic pipelines. MF and DCT wrote the manuscript, and other authors provided editorial input.

