## Supplementary figures and images for "Scaling up by looking closer: extending the range of moss-Nostoc symbioses deeper into central Europe and to new moss hosts in the forest canopy"

### Supplemental figure 1

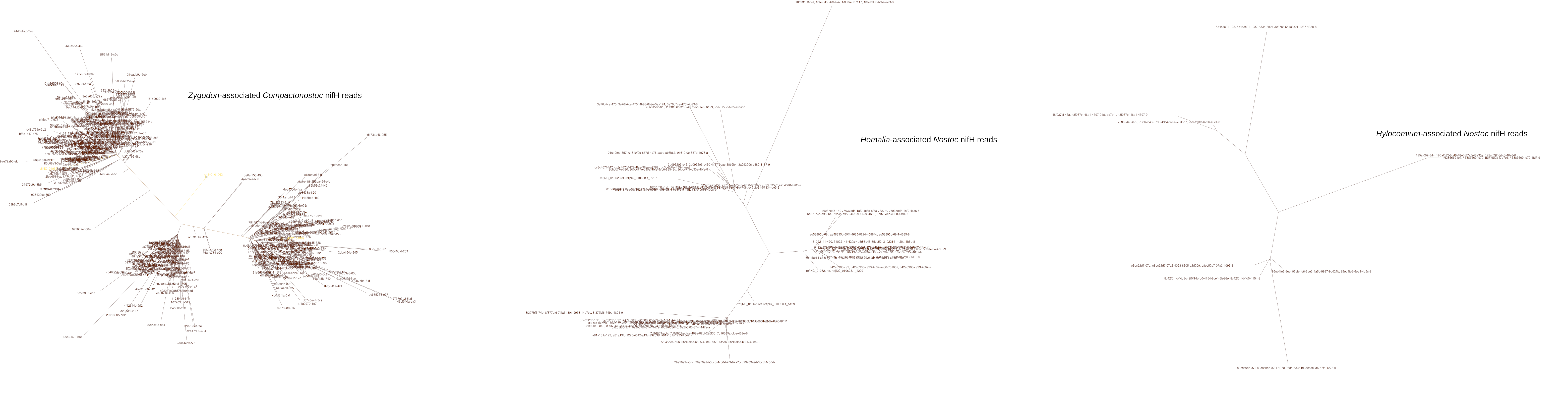

### Supplemental figure 2

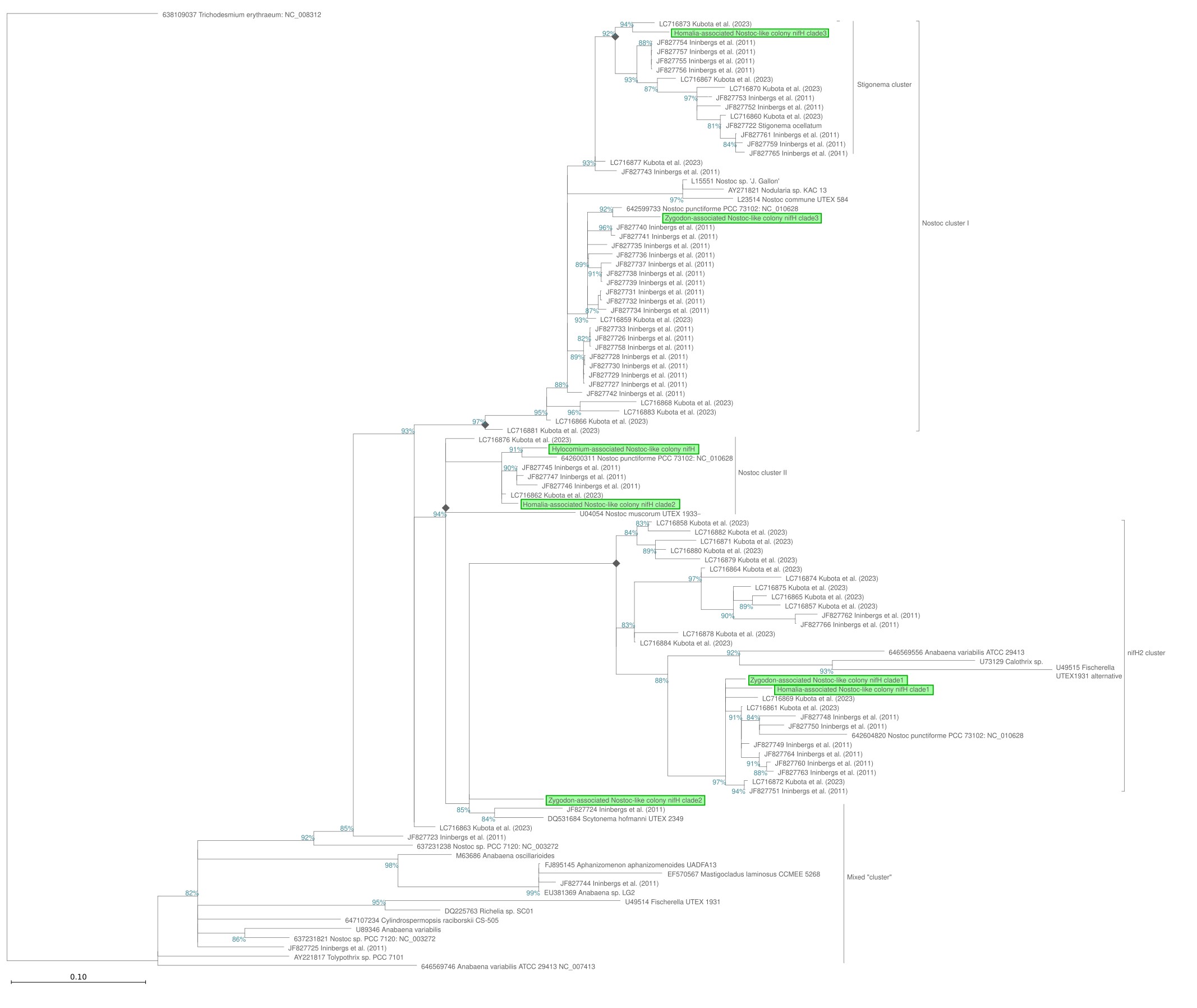
